# Multimodal Brain Age Gap as a Mediating Indicator in the Relation between Modifiable Dementia Risk Factors and Cognitive Functioning

**DOI:** 10.1101/2020.09.23.309369

**Authors:** Chang-Le Chen, Pin-Yu Chen, Yu-Hung Tung, Yung-Chin Hsu, Wen-Yih Isaac Tseng

## Abstract

**Introduction:** As a structural proxy for evaluating brain health, neuroimaging-based brain age gap (BAG) is presumed to link the dementia risks to cognitive changes in the premorbid phase, but this remains unclear.

**Methods:** Brain age prediction models were constructed and applied to a population-based cohort (N=371) to estimate their BAG. Further, structural equation modeling was employed to investigate the mediation effect of BAG between risk levels (assessed by 2 dementia-related risk scores) and cognitive changes (examined by 4 cognitive assessments).

**Results:** A higher burden of modifiable dementia risk factors was causally associated with a greater cognitive decline, and this was significantly mediated (*P*=0.017) by a larger multimodal BAG, which indicated an older brain. Moreover, a steeper slope (*P*=0.020) of association between cognitive decline and multimodal BAG was observed when individuals had higher dementia risks.

**Discussion:** Multimodal BAG is a potential mediating indicator to reflect the changes in the pathophysiological mechanism of cognitive aging.

## 1 INTRODUCTION

Brain aging is associated with compromised physical and mental condition [1] and increased risk of neurodegenerative diseases such as dementia [2]. The diversity in neurodegeneration suggests that the effect of aging on brain structure may be heterogeneous among individuals [3,4]. Improved understanding of the neural substrates underlying brain aging, therefore, is crucial to clarifying the heterochronicity of an aging brain. Developing a reliable marker for quantifying changes in brain structure is the first step in this process. A promising approach to identifying individual differences in brain aging is the exploitation of neuroimaging data for the estimation of brain age, which is regarded as a proxy of biological age of an individual’s brain [5].

A neuroimaging-based brain age is an imaging marker used to quantify the brain’s aging status [6]. With the aid of machine learning techniques, the brain scans acquired from healthy individuals are used to fit their chronological age to establish brain age prediction models, which can further be employed to predict other individuals’ brain age [7]. The discrepancy between an individual’s brain age and chronological age is called brain age gap (BAG), and it quantitatively indicates deviation from the defined healthy aging; higher BAG indicates an older brain [8]. Brain age measures have demonstrated extraordinary potential in detecting atypical brain aging [9-11] and, more crucially, in exhibiting associations with modifiable dementia-related risk factors [12,13] and cognitive changes [12,14].

Heterochronicity in cognitive aging can be attributed to modifiable factors such as lifestyle [12,15]. It has been postulated that lifestyle factors may affect brain structures, and these structural alterations then further affect cognitive functions [16,17]. However, whether BAG can serve as a mediating indicator in the cascaded causal association of the lifestyle–brain–cognition relationship remains unclear. It has been reported that higher cardiovascular risk factors may increase the incidence of multiple brain lesions and consequently lead to cognitive decline [18]. Thus, we hypothesize that BAG plays a mediating role in this causal relationship.

In this study, we examined the mediating effect of BAG on the causal association between modifiable risk factors and cognitive functioning in a premorbid population-based cohort. Specifically, we used multimodal neuroimaging data combined with a machine learning approach to estimate the BAG of gray matter (GM) and white matter (WM). Modifiable risk factors were measured using the well-validated Cardiovascular Risk Factors, Aging, and Incidence of Dementia (CAIDE) risk score [19] and the Framingham General Cardiovascular Risk Score (FGCRS) [20]. Cognitive functioning was defined by estimating the latent variable underlying 4 types of cognitive measurement [15]. Structural equation modeling (SEM) was employed to test the hypothetical mediation effect of BAG on the relationship between modifiable risk factors and cognitive functioning. To investigate the modulation of modifiable risk factors on the relation between BAG and cognition, we also explored the associations between BAG and cognitive functioning given different levels of risk factors.

## 2 METHODS

### 2.1 Participants

This retrospective study used the population-based data acquired by the Cambridge Centre for Ageing and Neuroscience (Cam-CAN) project (www.cam-can.com). The details of the recruitment protocol and ethical approval have been described in the studies of Shafto et al. [21] and Taylor et al. [22]. Briefly, after providing informed consent, all participants completed various sets of questionnaires, physical measurements, and cognitive examinations and received magnetic resonance imaging (MRI) scans. Participants were excluded if they had contraindications for MRI examination or a diagnosis of neurodegenerative diseases, psychiatric disorders, and other major health concerns [21]. A total of 636 premorbid participants aged 18–88 years were initially recruited in the study.

### 2.2 Image Analysis

#### 2.2.1 MRI Acquisition

MRI scanning was conducted using a 3T Siemens TIM Trio scanner with a 32-channel phased-array head coil [22]. High-resolution T1-weighted images were acquired using a three-dimensional magnetization-prepared rapid gradient echo sequence with voxel size = 1 mm isotropic. Two-shell diffusion tensor imaging (DTI) datasets were acquired using a twice-refocused diffusion pulsed-gradient spin-echo echo-planar imaging sequence, containing 30 directions with b = 1000 s/mm^2^, 30 directions with b = 2000 s/mm^2^, and 3 images with b = 0 s/mm^2^. Details of the imaging parameters are provided in Supplementary Information (SI)-S2.1.

#### 2.2.2 Image Feature Extraction

Before conducting image analyses, all T1-weighted images and DTI datasets underwent quality assurance (QA) procedures to ensure acceptable image quality. The detailed QA procedures are described in SI-S2.2. After this step, we had 616 participants aged 18–88 years for further analyses.

To extract GM features from T1-weighted images, voxel-based morphometry and surface-based morphometry were performed using Computational Anatomy Toolbox [23], an extension of Statistical Parametric Mapping Software [24] (SI-S2.3). Voxel-based morphometry [25] estimates voxel-wise GM volumes according to the LONI probabilistic whole-brain atlas, which contains 56 regions of interest (ROIs) [26]. Surface-based morphometry measures cortical thickness by using projection-based thickness estimation [27]. The Desikan–Killiany cortical atlas, which contains 68 cortical ROIs, was employed to sample cortical features [28]. In this manner, a total of 56 volumetric features and 68 cortical thickness features were obtained for estimating the GM-based BAG.

WM features were extracted from DTI datasets using an in-house analytic pipeline to automatically transform DTI data into tract-specific features [29] (SI-S2.3). Briefly, the DTI data were reconstructed, using the regularized version of mean apparent propagator MRI algorithm [30,31], into structure-related diffusion indices, namely the generalized fractional anisotropy (GFA) and mean diffusivity (MD). Next, a two-step registration with an advanced diffusion MRI-specific registration algorithm [32] was employed to minimize the registration bias arising from cross-lifespan data variation. Finally, the predefined tract coordinates on a standard template were projected, according to the transformation map from the registration process, onto individuals’ diffusion indices to sample tract-specific features. The pipeline produced 45 tract features for each index from each participant. Consequently, 45 GFA and 45 MD features were obtained to estimate the WM-based BAG.

### 2.3 Brain Age Modeling and Brain Age Gap Estimation

To estimate participants’ BAG from the Cam-CAN datasets, we fine-tuned the GM- and WM-based brain age pretrained models by using transfer learning [33]. First, the pretrained models, which employed a cascade neural network architecture, were established using the brain scans of a cognitively healthy cohort (N = 482) acquired at National Taiwan University Hospital (NTUH). Details of the NTUH cohort and the analytic procedure are provided in SI-S1.1. Next, the Cam-CAN cohort was split into tuning (N = 200), validation (N = 45), and target (N = 371) sets by using a conditional random method to ensure statistically identical age and sex distributions among the groups. These 3 sets were used to fine-tune the pretrained models, validate model performance, and estimate BAG for statistical analyses, respectively (SI-S1.2). After confirming the model performance, we applied the transferred GM- and WM-based brain age models to the target set to estimate their BAG (i.e., subtracting chronological age from the estimated brain age). Due to the age-related bias existing in the BAG [8], we removed the age effect from BAG before conducting further statistical analyses (SI-S4). Additionally, we provided biological inference of brain age models in SI-S5 for extra information and interpretation.

### 2.4 Assessment of Dementia Risk Burden

We assessed dementia risk burden by adopting the CAIDE and FGCRS scores, which have demonstrated clinical associations with cerebrovascular events, cognitive dysfunction, and brain pathology [18,34]. The CAIDE was the first validated tool for estimating dementia incidence risk and is widely used [35]. We adapted the formula of the CAIDE risk score by focusing on the modifiable items, which included education years, systolic blood pressure, body mass index, total cholesterol, and physical activity [36]. The FGCRS is used to evaluate cardiovascular burden [37] and is reportedly able to estimate dementia-related cognitive decline [34]. Likewise, we modified the formula of the FGCRS by evaluating the modifiable items, which included total cholesterol, systolic blood pressure measured with or without antihypertensive medication, smoking, diabetes, and cardiovascular history. Also, the FGCRS score was calculated according to the scoring standard for women. All participants in the target group had complete risk factor assessment data.

#### 2.5 Assessment of Cognitive Functions

General cognitive functioning was assessed using 4 well-known questionnaires. Specifically, we employed the Mini-Mental State Examination (MMSE) [38], Cattell’s Culture Fair Test of Fluid Intelligence, Scale II (CCFT) [39], Addenbrooke’s Cognitive Examination Revised version (ACE-R) [40], and the delayed phase of logical memory from the Wechsler Memory Scale Third UK Edition (WMS-D) [41] to measure participants’ cognitive abilities. The MMSE is a brief measure of cognitive functioning that has been widely used in clinical evaluation and research for screening patients with dementia [38]. The CCFT involves a general cognitive test that evaluates fluid intelligence [42]. The ACE-R, which measures multiple domain-specific cognition, can serve as a valid dementia screening test and is sensitive to early cognitive impairment [40]. The WMS-D is the delayed recall in logical memory test of the WMS [41], which has been shown to effectively reflect early deficits in memory functions [43]. We adopted these 4 cognitive scores to represent an individual’s cognitive functioning. All participants in the target group had complete cognitive data.

#### 2.6 Statistical Analyses

To investigate the causal association between modifiable risk factors, BAG, and cognitive functioning independent of unmodifiable factors, we conducted statistical bias adjustment to control for the confounding effect of age and sex on all the observed variables by using multiple linear regression. The residual of the estimated linear regression was used to represent the original observed variable for further analyses (SI-S4).

We employed SEM to test the mediating effect of BAG on the association between modifiable risk factors and cognitive functioning. In this study, 3 types of BAG were modeled, namely GM, WM, and multimodal BAG. Multimodal BAG was a latent variable representing global brain health and was synthesized from the GM- and WM-based BAG by using confirmatory factor analysis. Similarly, we estimated the latent variable “modifiable risk factor” using the CAIDE and FGCRS scores and the latent variable “cognitive functioning” using the MMSE, CCFT, ACE-R, and WMS-D scores. Mediation analysis was conducted to test the direct and indirect effects between the modifiable risk factor and the cognitive functioning. To assess the overall fit of SEM models, we used the comparative fit index (CFI) (good fit>0.95), standardized root-mean-square residual (SRMR) (good fit<0.08), and root-mean-squared error of approximation (RMSEA) (good fit<0.06). The estimation of SEM was performed using maximum likelihood with robust standard errors and mean- and variance-adjusted test statistics to account for non-normality. The *lavaan* package in R language was used for SEM analysis [44].

We also investigated the association between BAG and cognitive functioning under different levels of modifiable risk factors. We defined a group variable based on high- and low-risk factors according to the median split of the modifiable risk factor. For each type of BAG, a linear regression was fitted in which the dependent variable was the cognitive functioning and the independent variables were BAG, group variable for risk factor exposure, and their interaction. The Benjamini–Hochberg correction was used to address multiple comparison problem [45].

## 3 RESULTS

### 3.1 Participants’ Characteristics

Descriptive statistics for demographic, cognitive, and lifestyle variables are presented in Table 1. The tuning and validation groups were used to fine-tune the pretrained brain age models and evaluate model performance, respectively. With conditionally random split, the age distribution and sex proportion were statistically comparable between the tuning and validation groups (age: *t*_(243)_ = 1.152, *P* = 0.250; sex: χ^2^_(1)_ = 1.344, *P* = 0.246). The results of brain age modeling and model tuning with transfer learning were provided in SI-S3. Briefly, both the GM- and WM-based transferred models exhibited satisfactory performance in the Cam-CAN cohort. In the target group, the average BAG for GM and its SD were −0.04 (6.31) years, and the average BAG for WM and its SD were 0.51 (7.10) years. Additionally, we statistically removed the confounding effect of age and sex from the observed variables involving risk factors, cognitive scores, and BAG measures in the target group. The residuals of the observed variables, which were orthogonal to those confounding factors (*P* ≈ 1), were used for further analyses (SI-S4).

**Table 1.** Characteristics of the study participants

| Characteristics | Tuning group | Validation group | Target group |
| --- | --- | --- | --- |
| Sample size | 200 | 45 | 371 |
| Age (years) , mean (SD) | 55.7 (19.5) | 52.0 (19.3) | 53.7 (17.7) |
| Age range | 18-87 | 19-86 | 18-88 |
| Sex (female), % | 54.0% | 44.4% | 49.1% |
| ACE-R, mean (range) | 94.6 (76,100) | 94.7 (82,100) | 95.2 (76,100) |
| MMSE, mean (range) | 28.9 (25,30) | 29.0 (26,30) | 28.9 (25,30) |
| CCFT, mean (range) | 31.0 (13,43) | 32.1 (16,43) | 32.3 (12,44) |
| WMS-D, mean (range) | 12.6 (0,22) | 12.7 (4,21) | 13.1 (2,24) |
| CAIDE, mean (range) | 1.51 (0,7) | 1.38 (0,5) | 1.37 (0,9) |
| FGCRS, mean (range) | 1.52 (-3,14) | 1.07 (-3,14) | 1.61 (-3,14) |
| Education (years), mean (SD) | 14.1 (5.1) | 13.8 (4.7) | 14.5 (3.8) |
| Systolic pressure (mmHg), mean (SD) | 119.8 (16.7) | 120.4 (12.8) | 119.8 (17.0) |
| Body mass index (kg/m<sup>2</sup>), mean (SD) | 25.7 (4.4) | 24.9 (4.0) | 25.7 (4.4) |
| High cholesterol, n (%) | 31 (15.5%) | 7 (15.6%) | 43 (11.6%) |
| Physical activity (kJ/day/kg), mean (SD) | 40.8 (20.6) | 50.3 (34.8) | 44.2 (20.7) |
| Smoking, n (%) | 124 (62.0%) | 28 (62.2%) | 236 (63.6%) |
| Diabetes, n (%) | 9 (4.5%) | 2 (4.4%) | 15 (4.0%) |
| Antihypertensive medication, n (%) | 32 (16%) | 7 (15.6%) | 53 (14.3%) |
| Cardiovascular disease history, n (%) | 18 (9.0%) | 3 (6.7%) | 33 (8.9%) |

Abbreviations: SD: standard deviation; ACE-R: Addenbrooke's Cognitive Examination Revised version; MMSE: Mini-Mental State Examination; CCFT: Cattell's Culture Fair Test; WMS-D: delayed phase of logical memory from the Wechsler Memory Scale Third UK Edition; CAIDE: Cardiovascular Risk Factors, Aging, and Incidence of Dementia; FGCRS: Framingham General Cardiovascular Risk Score.

### 3.2 Structural Equation Modeling of Brain Age Gap

All BAG–mediated models exhibited favorable overall fitting in SEM. The fitting results were as follows: in the multimodal BAG–mediated model, CFI=0.954, SRMR=0.039, and RMSEA=0.053; in the GM-based BAG–mediated model, CFI=0.957, SRMR=0.036, and RMSEA=0.053; and in the WM-based BAG–mediated model, CFI=0.949, SRMR=0.038, and RMSEA=0.059.

Figure 1 displays the path diagrams of SEM. For the multimodal BAG–mediated model, the modifiable risk factors did not exhibit a significant direct effect on cognitive functioning (beta _(unstandardized)_ = −0.350, *P* = 0.150), but the indirect effect on cognitive functioning was significant for the multimodal BAG (beta_(unstandardized)_ = −0.229, *P* = 0.017), which indicated that the multimodal BAG had a significant mediation effect (percent mediation proportion: 39.6%) in the causal association between modifiable risk factors and cognitive functioning. For both the GM- and the WM-based BAG– mediated models, the modifiable risk factor had a significant direct effect on cognitive functioning (GM: beta_(unstandardized)_ = −0.457, *P* = 0.025; WM: beta_(unstandardized)_ = −0.504, *P* = 0.035), whereas the indirect effect through the GM- or the WM-based BAG on cognitive functioning was only marginally significant (GM: beta_(unstandardized)_ = −0.068, *P*=0.063; WM: beta_(unstandardized)_ = −0.079, *P* = 0.071), which indicated that the unimodal BAG (i.e., GM- or WM-based) had a marginally significant mediation effect (percent mediation proportion: GM: 13.0%; WM: 13.6%).

**Figure 1.**
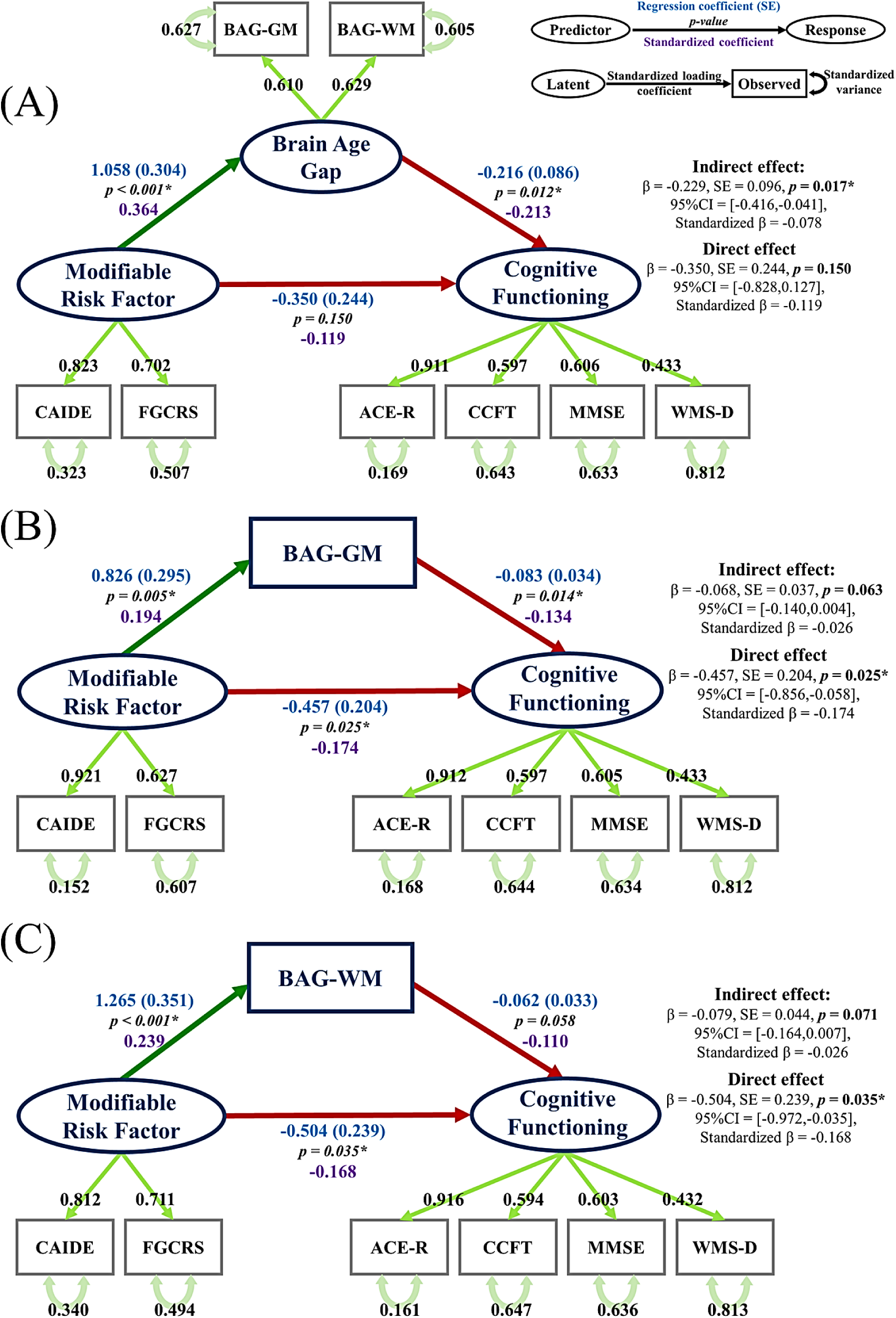
Path diagrams of structural equation models. Dark green arrows (denoting positive association) and red arrows (denoting negative association) are regression paths, starting from the predictor to the response. The statistics displayed beside the regression path are the regression coefficient and its standard error (blue), the corresponding *P* value (black), and the standardized coefficient (purple). The regression explanation is illustrated in the top-right corner of the figure. The ovals and squares indicate the latent variables and observed variables, respectively. The light green straight and curved arrows indicate the factor loading and error variance from the confirmatory factor analysis, respectively. Abbreviations: BAG: brain age gap; GM: gray matter; WM: white matter; CAIDE: Cardiovascular Risk Factors, Aging, and Incidence of Dementia risk score; FGCRS: Framingham General Cardiovascular Risk Score; ACE-R: Addenbrooke’s Cognitive Examination Revised version; Catt: Cattell’s Culture Fair Test of Fluid Intelligence, Scale II; MMSE: Mini-Mental State Examination; WMS-D: delayed phase of logical memory from the Wechsler Memory Scale Third UK Edition.

**Figure 2.**
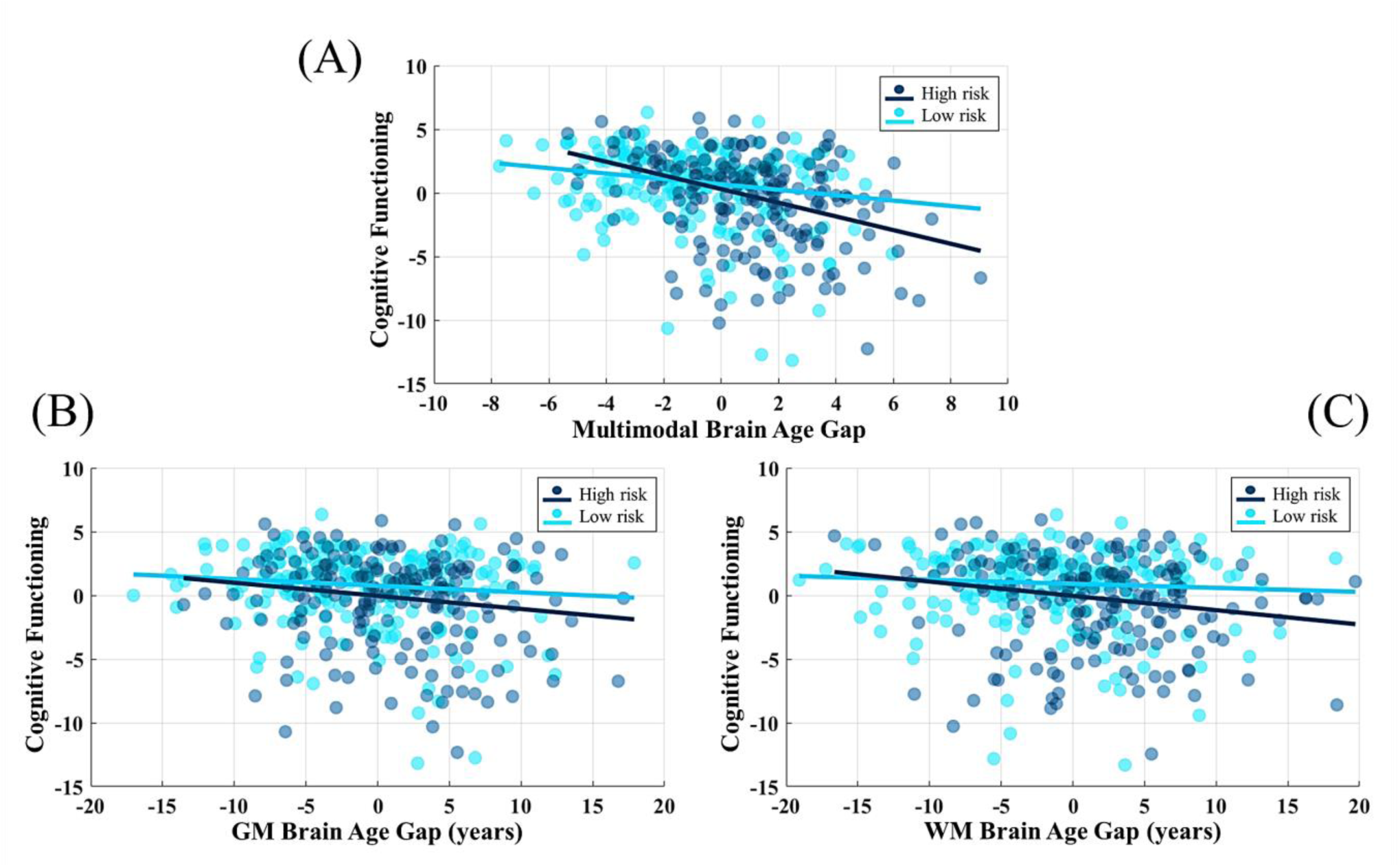
Regression plots of cognitive functioning relative to brain age gap under high- and low-risk modifiable factor levels. The solid lines indicate mean responses. The dark and light blue dots denote high- and low-risk groups, respectively.

In the regression paths between modifiable risk factors and BAG, multimodal BAG explained more of the variance (beta_(standardized)_ = 0.364, *P* < 0.001) in modifiable risk factors than the unimodal BAGs (GM: beta_(standardized)_ = 0.194, *P* = 0.005; WM: beta_(standardized)_ = 0.239, *P* < 0.001). Furthermore, all regression paths indicated that higher modifiable risk factors were significantly associated with larger BAG, which denoted premature aging status compared with the norm. In the regression paths between BAG and cognitive functioning, multimodal BAG also explained more variance in cognitive functioning (beta_(standardized)_ = −0.213, *P* = 0.012) than the other 2 measures (GM: beta_(standardized)_ = −0.134, *P* = 0.014; WM: beta_(standardized)_ = −0.110, *P* = 0.058). Larger multimodal and GM-based BAGs were significantly associated with lower cognitive functioning, whereas WM-based BAG exhibited only a marginal association with cognitive functioning.

### 3.3 Regression between Brain Age Gap and Cognitive Functioning under Different Risk Levels

After SEM analysis, we further stratified the target set into high- and low-risk-level groups by using a median split of the modifiable risk factor and investigated the associations between BAG and cognitive functioning (Fig.2). In the regression model of multimodal BAG, the BAG (beta = −0.216, *P* = 0.015) and its interaction with the risk level (beta = −0.322, *P* = 0.020) significantly explained cognitive functioning. This result indicated that, in the high-risk group, multimodal BAG had a greater negative slope for changes in cognitive functioning. However, in the regression models of unimodal BAG (i.e., GM and WM), the BAG (GM: beta = −0.055, *P* = 0.270; WM: beta = −0.034, *P* = 0.323)and its interaction with the risk level (beta = −0.047, *P* = 0.384; beta = −0.080, *P* = 0.212, respectively) did not significantly explain the dependent variables. Only the risk level was significant in the WM-based model (beta = −0.813, *P* = 0.041) and marginally significant in the GM-based model (beta = −0.735, *P* = 0.072), indicating that the high-risk group had inferior cognitive functioning relative to the low-risk group.

## 4 DISCUSSION

This study demonstrated that the neuroimaging-based BAG can serve as a mediating indicator between dementia-related modifiable risk factors and cognitive functioning. We obtained 2 major findings from the population-based cohort. First, the modifiable risk factor was causally associated with the cognitive functioning through the mediation of multimodal BAG. Second, under exposure to higher risk factors, multimodal BAG was associated with a greater declining slope in cognitive functioning.

Mediation testing indicated that the multimodal BAG represented a significant statistical link between modifiable risk factors and cognitive functioning; this implies that multimodal BAG, which employs both GM and WM information, more capably explained the variance in modifiable risk factors and the link with cognitive changes. A previous study revealed that mixed brain lesions were the structural mediator between cardiovascular risks and MMSE [18]. In our study, we further verified that the multimodal BAG, which condenses the information of major brain structures into a concise brain age index, can serve as a structural mediator between risk factors and cognition. Our results support the notion proposed in the theory of normal cognitive aging that modifiable risk factors shape the integrity of brain structures and consequentially change cognitive outcomes [16].

Cognitive aging may undergo different trajectories among individuals given diverse environmental exposure; the modifiable dementia-related risk factors, such as cardiovascular risks and lifestyle factors, substantially influence the cognitive aging process [16,46,47]. For instance, physical activity and educational attainment contribute to slowing cognitive aging [48], whereas vascular risks, such as hypertension and smoking, have an adverse effect and potentially accelerate cognitive decline [12,13]. In line with previous studies [16,18], our results revealed that the modifiable risk factor had a significant causal association with the cognitive functioning through either direct or indirect paths. Also, the significantly positive associations between the modifiable risk factor and all BAGs indicate that the brain structure ages prematurely under the conditions of higher modifiable risk factor. This suggests that the BAG based on either multimodal or unimodal features has sufficient sensitivity to reflect the levels of modifiable risk factor in a premorbid cohort. Additionally, we found that BAG was negatively correlated with the cognitive functioning, indicating that subtle changes in general cognition related to dementia can be captured by BAG, particularly when using multimodal features. This is consistent with the previous findings that BAG can reflect associations with several domain-specific cognitive measurements [12,14].

Furthermore, the strengths of the associations between multimodal BAG and cognitive functioning differed under distinct levels of the modifiable risk factor; the cognitive functioning declined more in the high-risk group given the same increment of BAG. This finding can support the life course model of the scaffolding theory of aging and cognition (STAC-R) [17]. The STAC-R is a conceptual model of cognitive aging that integrates evidence from structural and functional neuroimaging to explain how the combined effects of adverse and compensatory neural processes produce varying levels of cognitive function [17]. In the STAC-R model, life course experience leads to enrichment or depletion of neural resources. These changes in neural resources then affect brain structure, brain function, and compensatory scaffolding. Downstream, these three aspects further modulate the level of cognitive function. This theory posits that higher modifiable risk factors can decrease the enrichment and increase the depletion of neural resources. Once neural resources are exhausted, brain structures may deteriorate, and the compensatory scaffolding becomes more limited. These alterations consequently cause cognitive changes. We postulate that, given the same increment of BAG in the two risk groups, potentially because of the reduced scaffolding compensation, the high-risk group experienced more severe cognitive decline than did the low-risk group. It shows that multimodal BAG may be an underlying indicator of the structural alterations that connect the life course and cognition in the STAC-R model.

Brain age prediction models established using multiple imaging modalities provide valuable information on the brain’s aging process [8]. In this study, we initially created brain age models using different modality-specific features and then fine-tune the pretrained models to adapt to the Cam-CAN data by using transfer learning [33]. Our results revealed that transfer learning substantively improved model performance, particularly in the WM-based model, enabling the models to work accurately in a new site. Also, regional modeling of modality-specific brain aging patterns is capable of detecting associations with specific biomedical and clinical measures [12] and enables us to investigate biological inference of brain aging (SI-S5). Moreover, our results suggest that the multimodal BAG may be more sensitive than the unimodal BAG to reflect the structural changes in the lifestyle–brain– cognition relationship.

The strengths of this study include the use of a large population-based database with various measurements. A potential limitation of the present study is its cross-sectional design across lifespan; the cohort’s age differences may be a nuisance factor, which potentially confounds the study by overestimating the effect of interest. Hence, we removed the age effect from all observed variables throughout all statistical analyses. Nevertheless, a longitudinal study is warranted to verify and reproduce our results. Another limitation of this study is the lack of high-density lipoprotein data in the Cam-CAN project, which could have affected the definition of FGCRS. Finally, those enrolled in the present study were primarily from the cognitively normal population; the generalizability of the causal association in the lifestyle–brain–cognition relationship to other populations remains to be explored.

In summary, this population-based study demonstrates that an increased burden of modifiable risk factors contributes to multiple brain structure alterations, which can be quantified using a multimodal BAG measure. Higher BAG, indicating an older brain age compared with that of peers, is further associated with greater cognitive decline. Moreover, under greater exposure to the risk factors, an increase in BAG may accompany a greater decrease in cognition. Our findings in the premorbid population highlight that neuroimaging-based BAG can be a sensitive indicator to represent an individual’s brain aging status and is a validated mediating marker in the lifestyle–brain–cognition relationship.

## 6 Author’s Contribution

C.C. and W.T. conceived the study and were in charge of overall planning. P.C., Y.T., Y.H. helped in data collection of diffusion MRI datasets and verified the numerical results of the simulation. C.C. and Y.H. performed the image analyses. C.C. designed the prediction model, and conducted the experiments, statistical analyses and results visualization. C.C., P.C., Y.T. and W.T. contributed to the interpretation of the results. C.C. and W.T. wrote the manuscript. All authors discussed the results and commented on the manuscript.

### 7 Acknowledgement

This research was supported by National Health Research Institute (NHRI) Taiwan (grant: NHRI-EX109-10928NI). We are grateful to the data source of the Cam-CAN project (http://www.mrc-cbu.cam.ac.uk/datasets/camcan/). The Cam-CAN research project was supported by the Biotechnology and Biological Sciences Research Council (Grant #BB/H008217/1), and the European Union’s Horizon2020 research and innovation programme (Grant agreement #732592). We thank Wallace Academic Editing for assistance with English editing that greatly improved the manuscript.

## 9 Research data for this article

The data from Cambridge Centre for Ageing and Neuroscience (http://www.mrc-cbu.cam.ac.uk/datasets/camcan) are available online in accordance with their data access declaration. The data acquired from National Taiwan University Hospital (NTUH) are not available due to confidentiality agreement of NTUH Research Ethics Committee. Scripts of brain age modeling and transfer learning method are available in an online open-access repository (https://github.com/ChangleChen/BrainAge_TL). Code of imaging process conditionally available upon request from the corresponding author.

## 10 Ethical Approval

All procedures performed in this study involving human participants from the National Taiwan University Hospital (NTUH) were in accordance with the ethical standards of the NTUH Research Ethics Committee (REC) and with the 1964 Helsinki declaration and its later amendments or comparable ethical standards. Informed consent in the study was obtained from all individual participants who were recruited in the NTUH.

## Supplementary Information

S1 Brain Age Estimation S2 Image Data Processing

S3 Results of Brain Age Modeling and Model Tuning through Transfer Learning

S4 Statistical Bias Adjustment

S5 Biological Inference of Brain Age Models

References

## S1 Brain Age Estimation

### S1.1 Establishment of Brain Age Pretrained Models

The brain age models applied to the data from the Cambridge Centre for Ageing and Neuroscience (Cam-CAN) project were initially pretrained using the data from our National Taiwan University Hospital (NTUH) database. This database contained a training set (N = 482) and a test set (N = 70), which were used to establish brain age models and evaluate model performance, respectively (training set: mean age = 36.9 years, max = 92, min = 14, female proportion = 53.1%; test set: mean age = 36.8 years, max = 83, min = 14, female proportion = 52.2%). The distributions of age and sex in the 2 sets were statistically identical.

The neuroimaging data in the NTUH database were acquired using a 3T Siemens TIM Trio scanner with a 32-channel phased-array head coil, with the same imaging protocol used for all data collection. High-resolution T1-weighted imaging was performed using a 3D magnetization-prepared rapid gradient echo (3D-MPRAGE) sequence: repetition time/echo time (TR/TE) = 2000/3 ms, flip angle = 9°, field of view (FOV) = 256 × 192 × 208 mm^3^, and acquisition matrix = 256 × 192 × 208; this resulted in an isotropic spatial resolution of 1 mm^3^. The imaging protocol for diffusion-weighted images was the same as that for diffusion spectrum imaging (DSI). The DSI datasets were acquired using the diffusion pulsed-gradient spin-echo echo-planar imaging sequence with a twice-refocused balanced echo (1, 2): TR/TE = 9600/130 ms, slice thickness = 2.5 mm, acquisition matrix = 80 × 80, FOV = 200 × 200 mm^2^, and in-plane spatial resolution = 2.5 × 2.5 mm^2^. The diffusion-encoding acquisition scheme used in this dataset followed the DSI framework (2), in which were applied 102 diffusion-encoding gradients corresponding to the Cartesian grids in the half sphere of the 3D diffusion-encoding space within a radius of 3 units equivalent to b_max_ = 4000 s/mm^2^ (3). Because the data in the 3D diffusion-encoding space were real and symmetrical around the origin, the acquired half-sphere data were projected to fill the other half of the sphere.

The imaging feature processing was the same as that used for the Cam-CAN data. The details of the imaging process are described in S2. The input data for gray matter (GM)-based brain age modeling used the volume and cortical thickness features in regions of interest (ROIs), whereas that for white matter (WM)-based brain age modeling used the tract-specific features of generalized fractional anisotropy (GFA) and mean diffusivity (MD). The sex factor was also included as a predictor in the models. The neuroimaging features of the GM- and WM-based model input consisted of 124 and 90 features, respectively. Twelve-layer feed-forward cascade neural network models, which provide an accurate prediction with flexible model architecture for transfer learning, were used to predict age (4). The cascade neural network is a feed-forward neural network involving connections from the input and every previous layer to the subsequent layer. This network is similar to a simplified fully connected version of a dense block in densely connected convolutional networks, which avoid the vanishing-gradient problem and strengthen feature propagation (5). The loss function of model optimization was specified as a mean square error function, which was optimized using a gradient descent algorithm with an adaptive learning rate and constant momentum. A 10-fold cross-validation procedure was adopted within the training set to estimate brain age model performance. Validation set performance was used to stop the model parameter updates. The training procedure was implemented using MATLAB R2018b (MathWorks Inc., Natick, MA, USA) with an NVIDIA GeForce GTX 1080Ti (NVIDIA Inc., Santa Clara, CA, USA) graphics processing unit for accelerated computing. The performance of the trained brain age model was tested by predicting the brain age of individuals in the test set. To quantify model performance, Pearson’s correlation coefficient and the mean absolute error between the predicted age and chronological age were calculated. The results of brain age modeling were provided in Supplementary Information S3.

### S1.2 Fine-tune Pretrained Models to the Cam-CAN Datasets Using Transfer Learning

To transfer the GM- and WM-based brain age models from the NTUH cohort to the Cam-CAN dataset, the Cam-CAN cohort was split into tuning (N = 200), validation (N = 45), and target (N = 371) sets by using a conditional random method to ensure statistically identical age and sex distributions among the groups. These 3 sets were used to fine-tune the pretrained models, validate model performance, and estimate brain age gap for statistical analyses, respectively. According to our previous study (4), a sample size of approximately 200 observations as the tuning sample was able to produce a transferred model with satisfactory performance. The loss function of model optimization was specified as the mean square error function, which was optimized using a gradient descent algorithm with an adaptive learning rate and constant momentum. A 10-fold cross-validation procedure was conducted within the tuning set to estimate the performance of the transferred brain age model. The validation set was used to evaluate the model’s performance on an independent set. The modeling and tuning procedures were implemented using MATLAB R2018b (MathWorks Inc., Natick, MA, USA) with an NVIDIA GeForce GTX 1080Ti (NVIDIA Inc., Santa Clara, CA, USA) graphics processing unit for accelerated computing. Model performance was evaluated in terms of Pearson’s correlation coefficient (rho) and mean absolute error (MAE) between the estimated brain age and chronological age. The results of model tuning with transfer learning were provided in Supplementary Information S3. After confirming the model performance, we applied the transferred GM- and WM-based brain age models to the participants in the target set to estimate their brain age. We calculated brain age gap by subtracting chronological age from the estimated brain age.

## S2 Image Data Processing

### S2.1 MRI Imaging Parameters

The participants in the Cam-CAN dataset were scanned using a 3T Siemens TIM Trio scanner with a 32-channel phased-array head coil. High-resolution T1-weighted images were acquired using a 3D magnetization-prepared rapid gradient echo sequence: TR/TE = 2250/2.99 ms, inversion time = 900 ms, flip angle = 9°, FOV = 256 × 240 × 192 mm^3^, and voxel size = 1 mm isotropic. The diffusion-weighted images consisted of 2-shell diffusion tensor imaging (DTI) datasets, which were acquired using a twice-refocused diffusion pulsed-gradient spin-echo echo-planar imaging sequence with the following imaging parameters: TR = 9100 ms, TE = 104 ms; FOV = 192 × 192 mm^2^; voxel size = 2 mm isotropic; 66 axial slices using 30 directions with b = 1000 s/mm^2^, 30 directions with b = 2000 s/mm^2^, and 3 images with b = 0 s/mm^2^.

### S2.2 Image Quality Assurance

Before we performed data analysis, all T1-weighted images underwent quality assurance (QA) procedures, which are included in the Computational Anatomy Toolbox 12 (CAT12; http://dbm.neuro.uni-jena.de/cat.html), a novel retrospective QA framework for empirical quantification of quality differences. Retrospective QA involved automatic evaluation of essential image parameters such as noise, inhomogeneity, and image resolution. These quality measures were scaled to a rating scale, and “good” image quality level was required. Additional visual inspection was conducted to examine whether artifacts, including severe motion and abnormal lesions, remained in the images.

All diffusion datasets also underwent QA procedures, including examination of the signal-to-noise ratio (SNR), degree of alignment between T1- and diffusion-weighted images, and corrections for motion and eddy currents. The SNR was evaluated by calculating the mean signal of an object divided by the standard deviation (SD) of the background noise (6). In practice, the signal was determined using a central square of an image for each slice, and the noise was averaged from 4 corner regions. Diffusion datasets with an SNR higher than mean SNR minus 2.5 SDs at their site were included. The degree of within-subject alignment between T1- and diffusion-weighted images was evaluated by calculating the spatial correlation between the T1-weighted image–derived WM tissue probability map and the diffusion-weighted image–derived GFA map. Higher spatial correlation indicated greater spatial alignment between T1- and diffusion-weighted images. The datasets with coefficients of correlation higher than the mean spatial correlation minus 2.5 × SD were included. The correction algorithm for motion and eddy currents, EDDY in FSL (https://fsl.fmrib.ox.ac.uk/fsl/fslwiki/eddy), was used to detect and replace slices affected by signal loss due to bulk motion during diffusion encoding (7).

### S2.3 Image Feature Processing

In the image feature processing for GM, voxel-based morphometry and surface-based morphometry of 3D MPRAGE data were used. The image analyses were performed using an extension of the Statistical Parametric Mapping package (SPM12; Wellcome Department of Imaging Neuroscience, London, UK; www.fil.ion.ucl.ac.uk/spm) (8) called CAT12. For voxel-based morphometry analysis, the structural imaging data were preprocessed using the default settings of the CAT12 toolbox, including corrections for bias-field inhomogeneity and segmentation into GM, WM, and cerebrospinal fluid, followed by spatial normalization to the DARTEL (9) template in MNI space (voxel size: 1.5 × 1.5?×?1.5? mm^3^). Next, the LONI probabilistic brain atlas, containing 56 ROIs, was used as a reference of volumetric tissue compartments (10) to estimate the volume of each ROI. In addition, for surface-based morphometry analysis of cortical thickness, we applied the automated surface-preprocessing algorithms included in the CAT12 toolbox that enable the simultaneous estimation of cortical thickness of the left and right hemisphere by using the projection-based thickness method (11). Herein, cortical thickness was determined by estimating the WM distance based on tissue segmentation. We used WM distance and a derived neighbor relationship to project local maxima (which is equal to the cortical thickness) onto other GM voxels. This approach included partial volume correction and correction for sulcal blurring and sulcal asymmetries. Next, the Desikan–Killiany cortical atlas containing 68 cortical ROIs was employed to sample cortical features (12). In this manner, 56 volumetric features and 68 cortical thickness features were obtained to estimate GM-based brain age and calculate brain age gap.

In the image processing for WM, our in-house algorithm called tract-based automatic analysis (13) was employed. First, the diffusion indices, including GFA and MD, derived from the diffusion MRI dataset were computed using the regularization version of the framework of mean apparent propagator MRI (14, 15). The signal in 3D diffusion-encoding space was fitted with a series expansion of Hermite basis functions, which describe diffusion in various microstructural geometries (16). The zero-order term in the expansion series contained the diffusion tensor that characterizes the Gaussian displacement distribution. Higher-order terms in the expansion series were the orthogonal corrections to the Gaussian approximation, and these were used for reconstructing the average propagator. The MD in each voxel was determined by calculating the mean of the 3 eigenvalues of the diffusion tensor (17, 18). We quantified GFA as the SD of the orientation distribution function divided by the root mean square of the orientation distribution function (19). To extract effective features of WM, the diffusion indices were sampled according to the spatial coordinates of 45 predefined major fiber tract bundles over the whole brain, which were constructed in the DTI template NTU-DSI-122-DTI (20) through deterministic streamline-based tractography with multiple ROIs defined in the automated anatomical labeling atlas (21). In practice, the sampling coordinates were transformed from NTU-DSI-122-DTI into individual DTI datasets with the corresponding deformation maps. The deformation maps were obtained through 2-step registration, which included anatomical information provided by the T1-weighted images (22) and microstructural information provided by the DTI datasets (23). The sampling coordinates were aligned with the proceeding direction of each fiber tract bundle, and diffusion indices were sampled in the native space along the sampling coordinates normalized and divided into 100 steps. Furthermore, we averaged the indices along 100 steps. Finally, 45 GFA features and 45 MD features were obtained for estimating WM-based brain age and calculating brain age gap.

## S3 Results of Brain Age Modeling and Model Tuning through Transfer Learning

The GM- and WM-based brain age pretrained models were established using data from the NTUH database. Both the training set (N = 482) and independent test set (N = 70) resulted in a model with satisfactory performance in age prediction given the GM (training set: rho = 0.952, MAE = 4.36 years; test set: rho = 0.963, MAE = 4.73 years) and WM (training set: rho = 0.948, MAE = 4.59 years; test set: rho = 0.938, MAE = 5.04 years) features. These pretrained models were then fine-tuned using the tuning samples (N = 200) from the Cam-CAN database to fit the new data domain through transfer learning. Table 2 lists the prediction performance of the pretrained model and transferred model. The predictions in the Cam-CAN data directly made by the pretrained model were unacceptable, particularly in the WM-based brain age model. The MAE of the GM-based model was 7.87 years and 7.20 years for the validation set (N = 45) and target set (N = 371), respectively, and that of the WM-based model was 14.12 years and 14.93 years, respectively. After we performed transfer learning adjustment, both the GM- and WM-based brain age models exhibited acceptable performance in the independent validation set and the target set (GM-based model: validation set: MAE = 4.97 years; target set: 5.13 years; WM-based model: validation set: MAE = 5.38 years; target set: 5.56 years). The scores of brain age gap were further calculated by subtracting chronological age from the predicted age.

**Supplementary Table 1.**
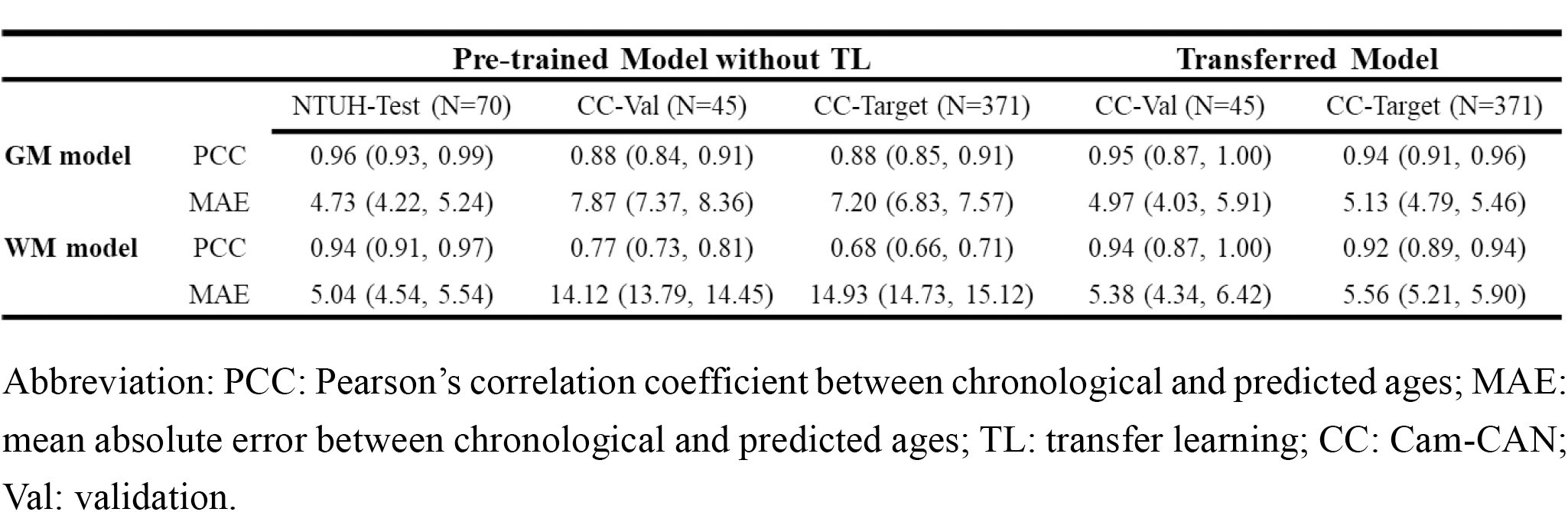
Prediction performance of the pretrained and transferred models

|  |  | Pre-trained Model without TL |  |  | Transferred Model |  |
| --- | --- | --- | --- | --- | --- | --- |
|  |  | NTUH-Test (N=70) | CC-Val (N=45) | CC-Target (N=371) | CC-Val (N=45) | CC-Target (N=371) |
| **GM model** | PCC | 0.96 (0.93, 0.99) | 0.88 (0.84, 0.91) | 0.88 (0.85, 0.91) | 0.95 (0.87, 1.00) | 0.94 (0.91, 0.96) |
|  | MAE | 4.73 (4.22, 5.24) | 7.87 (7.37, 8.36) | 7.20 (6.83, 7.57) | 4.97 (4.03, 5.91) | 5.13 (4.79, 5.46) |
| **WM model** | PCC | 0.94 (0.91, 0.97) | 0.77 (0.73, 0.81) | 0.68 (0.66, 0.71) | 0.94 (0.87, 1.00) | 0.92 (0.89, 0.94) |
|  | MAE | 5.04 (4.54, 5.54) | 14.12 (13.79, 14.45) | 14.93 (14.73, 15.12) | 5.38 (4.34, 6.42) | 5.56 (5.21, 5.90) |

Abbreviation: PCC: Pearson's correlation coefficient between chronological and predicted ages; MAE: mean absolute error between chronological and predicted ages; TL: transfer learning; CC: Cam-CAN; Val: validation.

## S4 Statistical Bias Adjustment

The observed variables involving risk factors, cognitive scores, and brain age gap had statistical dependency on age and sex, which could have had a nuisance effect. Thus, before statistical analysis in structural equation modeling, we conducted mass linear regression analysis to remove the confounding factors of age and sex from each observed variable. Specifically, in the multiple linear regression, the dependent variable was each observed variable of interest and the independent variables were age, age squared, and sex. The residual of the estimated linear regression was used to represent the original observed variable for further structural equation modeling analysis. The following table lists the statistical results of mass linear regression.

**Supplementary Table 2.** Results of bias adjustment

|  | Original Observed Variables |  | Adjusted Residual Variables |  |
| --- | --- | --- | --- | --- |
|  | Correlation with Age | Difference in Sex | Correlation with Age | Difference in Sex |
| ACE-R | rho = -0.368<br>(*p* = 0.000) | *t*-stat = -2.307<br>(*p* = 0.022) | rho = 0.000<br>(*p* = 1.000) | *t*-stat = 0.000<br>(*p* = 1.000) |
| CCFT | rho = -0.668<br>(*p* = 0.000) | *t*-stat = 0.918<br>(*p* = 0.359) | rho = 0.000<br>(*p* = 1.000) | *t*-stat = 0.000<br>(*p* = 1.000) |
| MMSE | rho = -0.363<br>(*p* = 0.000) | *t*-stat = -1.354<br>(*p* = 0.177) | rho = 0.000<br>(*p* = 1.000) | *t*-stat = 0.000<br>(*p* = 1.000) |
| WMS-D | rho = -0.414<br>(*p* = 0.000) | *t*-stat = -2.721<br>(*p* = 0.007) | rho = 0.000<br>(*p* = 1.000) | *t*-stat = 0.000<br>(*p* = 1.000) |
| CAIDE | rho = 0.485<br>(*p* = 0.000) | *t*-stat = 0.990<br>(*p* = 0.323) | rho = 0.000<br>(*p* = 1.000) | *t*-stat = 0.000<br>(*p* = 1.000) |
| FGCRS | rho = 0.472<br>(*p* = 0.000) | *t*-stat = 3.754<br>(*p* = 0.000) | rho = 0.000<br>(*p* = 1.000) | *t*-stat = 0.000<br>(*p* = 1.000) |
| BAG-GM | rho = -0.303<br>(*p* = 0.000) | *t*-stat = -0.965<br>(*p* = 0.335) | rho = 0.000<br>(*p* = 1.000) | *t*-stat = 0.000<br>(*p* = 1.000) |
| BAG-WM | rho = -0.336<br>(*p* = 0.000) | *t*-stat = -3.419<br>(*p* = 0.001) | rho = 0.000<br>(*p* = 1.000) | *t*-stat = 0.000<br>(*p* = 1.000) |

Note: Controlling for age, age square, and sex in the adjusted residual variables

## S5 Biological Inference of Brain Age Models

In addition to calculating brain age gap by using the brain age prediction model, we obtained information from the weights in the prediction models regarding the effects of the aging process on different brain features. To unveil this useful information underlying the neural network model, the permutation importance method (24) was used to calculate the importance of each imaging feature in the GM- and WM-based models separately. This method entailed repeated shuffling of the values of one predictor at a time, followed by making predictions using the resulting dataset. By comparing the predicted values with the true target values, we determined how much the shuffling affected the loss function because the importance of the variable shuffled was proportional to the performance drop. Finally, we normalized the scores of the loss function across all features from 0 to 1 to enable direct comparison.

In the result of feature contributions to brain age estimation, we adopted the proportional contribution of the features from the top 50% of importance and normalized them into percent ratios to represent contribution rates. Supplementary Figure 1 illustrates the feature importance derived from the brain age models. Given the measurements of feature type in GM, regional volume (51.3% contribution rate) had an effect comparable to that of cortical thickness (48.7%). GFA values had a higher contribution rate (54.2%) than did MD values (45.8%). In terms of hemispheric regions, features in the left and right hemispheres had comparable aging effects on both GM (left: 48.7%, right: 51.3%) and WM (left: 49.1%, right: 50.9%). We further divided the GM features into 5 major anatomical regions: frontal, parietal, occipital, temporal, and limbic and other regions. The temporal lobes (27.7%), limbic and others (22.0%), and frontal regions (20.6%) contributed the most, followed by the parietal (15.1%) and occipital (14.5%) regions. The WM features were partitioned into the association, projection, and corpus callosum fiber systems. The corpus callosum represented almost half of the contribution (48.4%) in the WM features, followed by the association fiber system (31.2%) and the projection fiber system (20.4%).

To our knowledge, estimation of brain age from multidimensional neuroimaging features typically relies on machine learning or deep learning techniques to perform the regression analysis. These advanced modeling approaches, particularly deep learning, are regarded as “black boxes” because the models themselves are generally infrequently interpretable. We used the permutation importance calculation method to assess the interpretability of the brain age model. In the GM-based model, the contributions of cortical volume and cortical thickness were relatively comparable, suggesting that these 2 measures are equally essential in capturing the changes in GM morphology during the aging process (25, 26). Nevertheless, the regional contributions were distinct across major anatomical lobes: the temporal, frontal, and limbic regions and others involving the basal ganglia, brain stem, and cerebellum had higher averaged weights than the parietal and occipital regions, consistent with age-related alterations in the brain reported previously (26, 27). Furthermore, although a related study reported that MD is a more age-sensitive diffusion measure than other indices (28), in our WM-based model, GFA had a higher contribution to brain age prediction than did MD, suggesting that GFA may be more suitable than MD in reflecting the microstructural changes of the brain over an individual’s lifespan. As for the 3 major fiber systems, callosal fibers contributed the most, followed by association fibers and project fibers, which is consistent with the aging patterns of the WM reported in relevant studies (28, 29). Regarding the difference between bilateral hemispheres, although regional aging lateralization has been demonstrated in several studies (30, 31), no apparent lateralization between bilateral hemispheres was found in the present study. Our feature importance estimation generally agreed with the observations reported in previous brain aging studies, suggesting that machine learning–based models automatically capture the aging features that are consistent with the current domain knowledge.

**Supplementary Figure 1.**
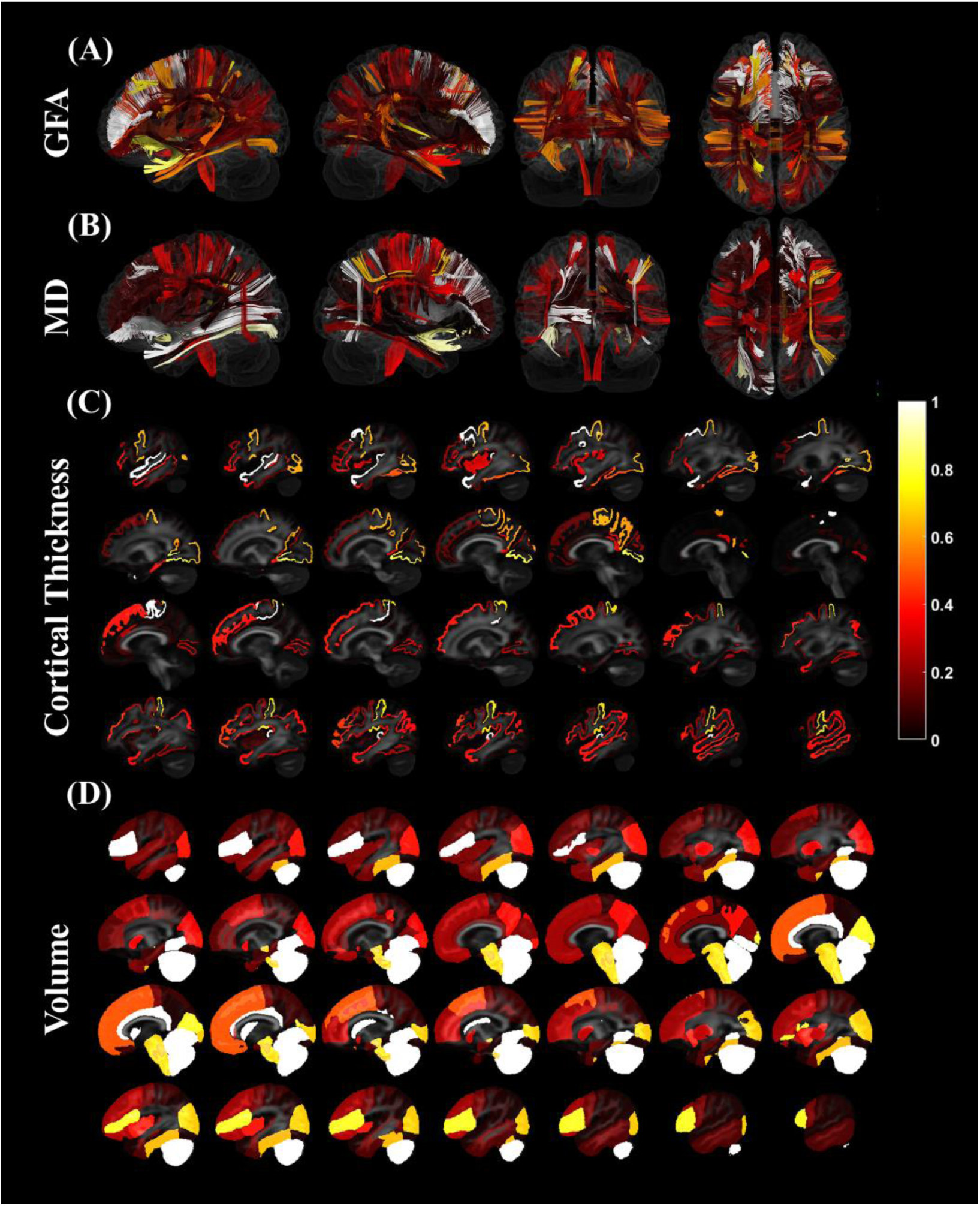
Salience map of feature importance in the brain age prediction model. The tract-specific features derived from generalized fractional anisotropy (GFA, subplot A) and mean diffusivity (MD, subplot B) contribute to the white matter brain age model. The region-specific features based on cortical thickness (C) and gray matter volume (D) contribute to the gray matter brain age model. Regional values closer to 1 indicate higher importance relative to the other features.

## References

[1] Cole JH, Ritchie SJ, Bastin ME, Hernández MV, Maniega SM, Royle N, et al. Brain age predicts mortality. Molecular psychiatry. 2018;23:1385–92.

[2] Franke K, Gaser C. Longitudinal changes in individual BrainAGE in healthy aging, mild cognitive impairment, and Alzheimer’s disease. GeroPsych: The Journal of Gerontopsychology and Geriatric Psychiatry. 2012;25:235.

[3] Deary IJ, Corley J, Gow AJ, Harris SE, Houlihan LM, Marioni RE, et al. Age-associated cognitive decline. British medical bulletin. 2009;92:135–52.

[4] Wyss-Coray T. Ageing, neurodegeneration and brain rejuvenation. Nature. 2016;539:180–6.

[5] Cole JH, Franke K. Predicting age using neuroimaging: innovative brain ageing biomarkers. Trends in neurosciences. 2017;40:681–90.

[6] Franke K, Gaser C. Ten years of BrainAGE as a neuroimaging biomarker of brain aging: What insights have we gained? Frontiers in neurology. 2019;10:789.

[7] Cole JH, Franke K. Predicting Age Using Neuroimaging: Innovative Brain Ageing Biomarkers. Trends Neurosci. 2017;40:681–90.

[8] Smith SM, Vidaurre D, Alfaro-Almagro F, Nichols TE, Miller KL. Estimation of brain age delta from brain imaging. Neuroimage. 2019;200:528–39.

[9] Gaser C, Franke K, Klöppel S, Koutsouleris N, Sauer H, Initiative AsDN. BrainAGE in mild cognitive impaired patients: predicting the conversion to Alzheimer’s disease. PloS one. 2013;8:e67346.

[10] Kaufmann T, van der Meer D, Doan NT, Schwarz E, Lund MJ, Agartz I, et al. Common brain disorders are associated with heritable patterns of apparent aging of the brain. Nature neuroscience. 2019;22:1617–23.

[11] Chen CL, Shih YC, Liou HH, Hsu YC, Lin FH, Tseng WI. Premature white matter aging in patients with right mesial temporal lobe epilepsy: A machine learning approach based on diffusion MRI data. Neuroimage Clin. 2019;24:102033.

[12] Cole JH. Multi-modality neuroimaging brain-age in UK Biobank: relationship to biomedical, lifestyle and cognitive factors. Neurobiology of Aging. 2020.

[13] de Lange A-MG, Anatürk M, Kaufmann T, Cole JH, Griffanti L, Zsoldos E, et al. Multimodal brain-age prediction and cardiovascular risk: The Whitehall II MRI sub-study. bioRxiv. 2020.

[14] Boyle R, Jollans L, Rueda-Delgado LM, Rizzo R, Yener GG, McMorrow JP, et al. Brain-predicted age difference score is related to specific cognitive functions: A multi-site replication analysis. Brain Imaging and Behavior. 2020:1–19.

[15] Borgeest GS, Henson RN, Shafto M, Samu D, Cam-CAN, Kievit RA. Greater lifestyle engagement is associated with better age-adjusted cognitive abilities. Plos one. 2020;15:e0230077.

[16] Harada CN, Love MCN, Triebel KL. Normal cognitive aging. Clinics in geriatric medicine. 2013;29:737–52.

[17] Reuter-Lorenz PA, Park DC. How does it STAC up? Revisiting the scaffolding theory of aging and cognition. Neuropsychology review. 2014;24:355–70.

[18] Wang R, Fratiglioni L, Kalpouzos G, Lövdén M, Laukka EJ, Bronge L, et al. Mixed brain lesions mediate the association between cardiovascular risk burden and cognitive decline in old age: A population-based study. Alzheimer’s & Dementia. 2017;13:247–56.

[19] Luchsinger JA, Mayeux R. Cardiovascular risk factors and Alzheimer’s disease. Current atherosclerosis reports. 2004;6:261–6.

[20] Kannel WB, McGee D, Gordon T. A general cardiovascular risk profile: the Framingham Study. The American journal of cardiology. 1976;38:46–51.

[21] Shafto MA, Tyler LK, Dixon M, Taylor JR, Rowe JB, Cusack R, et al. The Cambridge Centre for Ageing and Neuroscience (Cam-CAN) study protocol: a cross-sectional, lifespan, multidisciplinary examination of healthy cognitive ageing. BMC neurology. 2014;14:204.

[22] Taylor JR, Williams N, Cusack R, Auer T, Shafto MA, Dixon M, et al. The Cambridge Centre for Ageing and Neuroscience (Cam-CAN) data repository: Structural and functional MRI, MEG, and cognitive data from a cross-sectional adult lifespan sample. Neuroimage. 2017;144:262–9.

[23] Gaser C, Dahnke R. CAT-a computational anatomy toolbox for the analysis of structural MRI data. HBM. 2016;2016:336–48.

[24] Ashburner J, Barnes G, Chen C, Daunizeau J, Flandin G, Friston K. SPM12 Manual. Wellcome Trust Centre for Neuroimaging, London. 2014.

[25] Ashburner J, Friston KJ. Voxel-based morphometry—the methods. Neuroimage. 2000;11:805–21.

[26] Shattuck DW, Mirza M, Adisetiyo V, Hojatkashani C, Salamon G, Narr KL, et al. Construction of a 3D probabilistic atlas of human cortical structures. Neuroimage. 2008;39:1064–80.

[27] Dahnke R, Yotter RA, Gaser C. Cortical thickness and central surface estimation. Neuroimage. 2013;65:336–48.

[28] Desikan RS, Ségonne F, Fischl B, Quinn BT, Dickerson BC, Blacker D, et al. An automated labeling system for subdividing the human cerebral cortex on MRI scans into gyral based regions of interest. Neuroimage. 2006;31:968–80.

[29] Chen YJ, Lo YC, Hsu YC, Fan CC, Hwang TJ, Liu CM, et al. Automatic whole brain tract-based analysis using predefined tracts in a diffusion spectrum imaging template and an accurate registration strategy. Human brain mapping. 2015;36:3441–58.

[30] Özarslan E, Koay CG, Shepherd TM, Komlosh ME, Irfanoglu MO, Pierpaoli C, et al. Mean apparent propagator (MAP) MRI: a novel diffusion imaging method for mapping tissue microstructure. NeuroImage. 2013;78:16–32.

[31] Hsu YC, Tseng WY. An efficient regularization method for diffusion MAP-MRI estimation. 2018 ISMRM-ESMRMB Joint Annual Meeting. 2018.

[32] Hsu YC, Hsu CH, Tseng WY. A large deformation diffeomorphic metric mapping solution for diffusion spectrum imaging datasets. Neuroimage. 2012;63:818–34.

[33] Chen C-L, Hsu Y-C, Yang L-Y, Tung Y-H, Luo W-B, Liu CM, et al. Generalization of diffusion magnetic resonance imaging–based brain age prediction model through transfer learning. NeuroImage. 2020:116831.

[34] Kaffashian S, Dugravot A, Elbaz A, Shipley MJ, Sabia S, Kivimäki M, et al. Predicting cognitive decline: A dementia risk score vs the Framingham vascular risk scores. Neurology. 2013;80:1300–6.

[35] Sindi S, Calov E, Fokkens J, Ngandu T, Soininen H, Tuomilehto J, et al. The CAIDE Dementia Risk Score App: The development of an evidence-based mobile application to predict the risk of dementia. Alzheimer’s & Dementia: Diagnosis, Assessment & Disease Monitoring. 2015;1:328–33.

[36] Kivipelto M, Ngandu T, Laatikainen T, Winblad B, Soininen H, Tuomilehto J. Risk score for the prediction of dementia risk in 20 years among middle aged people: a longitudinal, population-based study. The Lancet Neurology. 2006;5:735–41.

[37] Kaffashian S, Dugravot A, Nabi H, Batty GD, Brunner E, Kivimäki M, et al. Predictive utility of the Framingham general cardiovascular disease risk profile for cognitive function: evidence from the Whitehall II study. European heart journal. 2011;32:2326–32.

[38] Folstein MF, Folstein SE, McHugh PR. “Mini-mental state”: a practical method for grading the cognitive state of patients for the clinician. Journal of psychiatric research. 1975;12:189–98.

[39] Cattell RB. Abilities: Their structure, growth, and action. 1971.

[40] Mioshi E, Dawson K, Mitchell J, Arnold R, Hodges JR. The Addenbrooke’s Cognitive Examination Revised (ACE-R): a brief cognitive test battery for dementia screening. International Journal of Geriatric Psychiatry: A journal of the psychiatry of late life and allied sciences. 2006;21:1078–85.

[41] Wechsler D. Wechsler Memory Scale Third UK E. London: Harcourt Assessment; 1999.

[42] Tucker-Drob EM. Cognitive Aging and Dementia: A Life-Span Perspective. Annual Review of Developmental Psychology. 2019;1:177–96.

[43] Johnson DK, Storandt M, Balota DA. Discourse analysis of logical memory recall in normal aging and in dementia of the Alzheimer type. Neuropsychology. 2003;17:82.

[44] Rosseel Y. Lavaan: An R package for structural equation modeling and more. Version 0.5–12 (BETA). Journal of statistical software. 2012;48:1–36.

[45] Benjamini Y, Hochberg Y. Controlling the false discovery rate: a practical and powerful approach to multiple testing. Journal of the Royal statistical society: series B (Methodological). 1995;57:289–300.

[46] Kramer AF, Bherer L, Colcombe SJ, Dong W, Greenough WT. Environmental influences on cognitive and brain plasticity during aging. The Journals of Gerontology Series A: Biological Sciences and Medical Sciences. 2004;59:M940–M57.

[47] Stern Y. Cognitive reserve in ageing and Alzheimer’s disease. The Lancet Neurology. 2012;11:1006–12.

[48] Steffener J, Habeck C, O’Shea D, Razlighi Q, Bherer L, Stern Y. Differences between chronological and brain age are related to education and self-reported physical activity. Neurobiology of aging. 2016;40:138–44.

## References

1. Reese, T.G., Heid, O., Weisskoff, R.M., Wedeen, V.J., 2003. Reduction of eddy-current-induced distortion in diffusion MRI using a twice-refocused spin echo. Magn. Reson. Med. 49, 177–182.

2. Wedeen, V.J., Hagmann, P., Tseng, W.Y., Reese, T.G., Weisskoff, R.M., 2005. Mapping complex tissue architecture with diffusion spectrum magnetic resonance imaging. Magn. Reson. Med. 54, 1377–1386.

3. Kuo, L.W., Chen, J.H., Wedeen, V.J., Tseng, W.Y., 2008. Optimization of diffusion spectrum imaging and q-ball imaging on clinical MRI system. Neuroimage. 41, 7–18.

4. Chen CL, et al. (2020) Generalization of diffusion magnetic resonance imaging-based brain age prediction model through transfer learning. Neuroimage 217:116831.

5. Huang, G., Liu, Z., Van Der Maaten, L., Weinberger, K.Q., 2017. Densely connected convolutional networks, in Proceedings of the IEEE Conference on Computer Vision and Pattern Recognition. pp. 2261–2269.

6. Dietrich, O., Raya, J.G., Reeder, S.B., Reiser, M.F., Schoenberg, S.O., 2007. Measurement of signal-to-noise ratios in MR images: influence of multichannel coils, parallel imaging, and reconstruction filters. J. Magn. Reson. Imaging. 26, 375–385.

7. Andersson, J.L.R., Sotiropoulos, S.N., 2016. An integrated approach to correction for off-resonance effects and subject movement in diffusion MR imaging. Neuroimage. 125, 1063–1078.

8. Ashburner, J., Barnes, G., Chen, C., Daunizeau, J., Flandin, G., Friston, K., Kiebel, S., Kilner, J., Litvak, V., Moran, R., 2014. SPM12 manual. Wellcome Trust Centre for Neuroimaging, London, UK, 2464.

9. Ashburner, J., Friston, K.J., 2011. Diffeomorphic registration using geodesic shooting and Gauss-Newton optimisation. Neuroimage. 55, 954–967.

10. Shattuck, D.W., Mirza, M., Adisetiyo, V., Hojatkashani, C., Salamon, G., Narr, K.L., Poldrack, R.A., Bilder, R.M., Toga, A.W., 2008. Construction of a 3D probabilistic atlas of human cortical structures. Neuroimage 39, 1064–1080.

11. Dahnke, R., Ziegler, G., Gaser, C., 2012. Local adaptive segmentation. Beijing. HBM. Available online at: http://dbm.neuro.uni-jena.de/HBM2012/HBM2012-Dahnke02.pdf.

12. Desikan, R.S., Ségonne, F., Fischl, B., Quinn, B.T., Dickerson, B.C., Blacker, D., Buckner, R.L., Dale, A.M., Maguire, R.P., Hyman, B.T., 2006. An automated labeling system for subdividing the human cerebral cortex on MRI scans into gyral based regions of interest. Neuroimage 31, 968–980.

13. Chen, Y.J., Lo, Y.C., Hsu, Y.C., Fan, C.C., Hwang, T.J., Liu, C.M., et al., 2015. Automatic whole brain tract-based analysis using predefined tracts in a diffusion spectrum imaging template and an accurate registration strategy. Hum. Brain Mapp. 36, 3441–3458.

14. Hsu, Y.C., Tseng, W.Y., 2018. An efficient regularization method for diffusion MAP-MRI estimation. 2018 ISMRM-ESMRMB Joint Annual Meeting, Paris, France.

15. Ozarslan, E., Koay, C.G., Shepherd, T.M., Komlosh, M.E., Irfanoglu, M.O., Pierpaoli, C., et al., 2013. Mean apparent propagator (MAP) MRI: a novel diffusion imaging method for mapping tissue microstructure. Neuroimage. 78, 16–32.

16. Avram, A.V., Sarlls, J.E., Barnett, A.S., Ozarslan, E., Thomas, C., Irfanoglu, M.O., et al., 2016. Clinical feasibility of using mean apparent propagator (MAP) MRI to characterize brain tissue microstructure. Neuroimage. 127, 422–434.

17. Alexander, A.L., Lee, J.E., Lazar, M., Field, A.S., 2007. Diffusion Tensor Imaging of the Brain. Neurotherapeutics. 4, 316–329.

18. Le Bihan, D., Mangin, J.F., Poupon, C., Clark, C.A., Pappata, S., Molko, N., et al., 2001. Diffusion tensor imaging: concepts and applications. J. Magn. Reson. Imaging. 13, 534–546.

19. Tuch, D.S., 2004. Q-ball imaging. Magn. Reson. Med. 52, 1358–1372.

20. Hsu, Y.C., Lo, Y.C., Chen, Y.J., Wedeen, V.J., Isaac Tseng, W.Y., 2015. NTU-DSI-122: A diffusion spectrum imaging template with high anatomical matching to the ICBM-152 space. Hum. Brain Mapp. 36, 3528–3541.

21. Tzourio-Mazoyer, N., Landeau, B., Papathanassiou, D., Crivello, F., Etard, O., Delcroix, N., et al., 2002. Automated anatomical labeling of activations in SPM using a macroscopic anatomical parcellation of the MNI MRI single-subject brain. Neuroimage. 15, 273–289.

22. Ashburner, J., Friston, K.J., 2000. Voxel-based morphometry—the methods. Neuroimage 11, 805–821.

23. Hsu, Y.-C., Hsu, C.-H., Tseng, W.-Y.I., 2012. A large deformation diffeomorphic metric mapping solution for diffusion spectrum imaging datasets. NeuroImage. 63, 818–834.

24. Altmann A, Toloşi L, Sander O, Lengauer T. Permutation importance: a corrected feature importance measure. Bioinformatics. 2010;26:1340–7.

25. Thambisetty M, Wan J, Carass A, An Y, Prince JL, Resnick SM. Longitudinal changes in cortical thickness associated with normal aging. Neuroimage. 2010;52:1215–23.

26. Alexander GE, Chen K, Merkley TL, Reiman EM, Caselli RJ, Aschenbrenner M, et al. Regional network of magnetic resonance imaging gray matter volume in healthy aging. Neuroreport. 2006;17:951–6.

27. Raji CA, Lopez OL, Kuller LH, Carmichael OT, Longstreth Jr WT, Gach HM, et al. White matter lesions and brain gray matter volume in cognitively normal elders. Neurobiology of aging. 2012;33:834. e7-. e16.

28. Cox SR, Ritchie SJ, Tucker-Drob EM, Liewald DC, Hagenaars SP, Davies G, et al. Ageing and brain white matter structure in 3,513 UK Biobank participants. Nature communications. 2016;7:1–13.

29. Yeatman JD, Wandell BA, Mezer AA. Lifespan maturation and degeneration of human brain white matter. Nature communications. 2014;5:1–12.

30. Guadalupe T, Mathias SR, Theo G, Whelan CD, Zwiers MP, Abe Y, et al. Human subcortical brain asymmetries in 15,847 people worldwide reveal effects of age and sex. Brain imaging and behavior. 2017;11:1497–514.

31. Kong X-Z, Mathias SR, Guadalupe T, Glahn DC, Franke B, Crivello F, et al. Mapping cortical brain asymmetry in 17,141 healthy individuals worldwide via the ENIGMA Consortium. Proceedings of the National Academy of Sciences. 2018;115:E5154–E63.

